# Visual working memory-related saccade biases are amplified by task demands, without updating working memory content

**DOI:** 10.1101/2025.10.24.684327

**Authors:** Patrik Polgári, Alexander C. Schütz

## Abstract

Visual working memory (VWM) and oculomotor control exert bi-directional influences on each other: eye movements during memory delays can influence VWM performance, and VWM content can bias the execution of eye movements. Oculomotor capture has been shown to extend to automatic, corrective saccades that can be biased towards distractors whose feature matches VWM content. We address the question of bi-directional influences between VWM and the oculomotor system, here with a focus on corrective saccade biases. 20 participants had to memorize a color hue and later discriminate it from another hue from a different (Easy condition) or the same color category (Difficult condition). Between encoding and test, they performed a gaze-contingent saccade task where, mid-saccade, the position of items shifted, introducing an artificial saccade error, and the color of a distractor neighboring the target changed to a color (mis)matching the memorized one. VWM performance was higher in the Easy compared to the Difficult experimental block, reflecting the successful manipulation of task demands. Corrective saccade biases towards the distractor occurred more frequently when it matched memory content compared to when it did not, replicating previous findings. Importantly, they also occurred more in the Difficult experimental block, which required stronger VWM representations than the Easy condition. VWM performance, however, was not affected by saccade corrections to memory (mis)matching distractors. Our results provided no evidence of saccade biases having a role in the updating of VWM content, however they indicate an influence of task demands and VWM engagement on automatic oculomotor biases.

## 1. Introduction

Vision is our primary mode of access to our environment, and the amount and quality of acquired visual information depends, due to anatomical restrictions, on scanning eye movements. The execution of saccades and fixations is, however, not enough for efficient perception. The oculomotor system shares an obligatory relationship not only with attentional networks (Schneider & Deubel, 1995; Deubel & Schneider, 1996; Deubel, 2008; Rolfs et al., 2011), but, as suggested by recent emerging evidence, also with visual working memory (VWM) (Van der Stigchel & Hollingworth, 2018).

Interestingly, VWM and eye movements exert bi-directional influences on each other (van Ede, 2020). Eye movements executed during memory delays can have beneficial (Hanning et al., 2016; Ohl & Rolfs, 2017, 2018; Hanning & Deubel, 2018), or detrimental effects (Lawrence et al., 2004; Schut et al., 2017) on VWM performance. Conversely, the content of VWM has been shown to influence oculomotor control (Hayhoe et al., 1998). This can be a repulsive effect, biasing saccade trajectories away from a remembered object’s location (Theeuwes et al., 2005; Belopolsky & Theeuwes, 2011; Boon et al., 2019), or an attractive effect, where a memorized item biases attention (“attentional capture”) and (micro-)saccades (“oculomotor capture”) towards similar featured objects (Downing & Dodds, 2004; Woodman & Luck, 2004; Soto et al., 2005, 2006; Olivers et al., 2006; Hollingworth et al., 2013b; Olmos-Solis et al., 2017; van Loon et al., 2017; Foerster & Schneider, 2018, 2020).

Hollingworth & Luck (2009) showed that oculomotor capture extends also to a more automatic form of eye movements: corrective saccades. This type of eye movement occurs often in visual search, when a saccade does not reach the exact target position, producing a saccade error (Frost & Pöppel, 1976; Kapoula, 1985; Hollingworth et al., 2008). This saccade error is swiftly corrected by the oculomotor system via a secondary saccade, bringing the gaze onto the originally intended target, in a way that observers are generally not aware of the correction. Hollingworth & Luck (2009) used a gaze-contingent paradigm to induce artificial saccade errors and manipulate the color of a distractor object neighboring a saccade target. Corrective saccades were biased away from the original saccade target and towards the distractor more often when the distractor changed to the color held in VWM, compared to a condition where it changed to an unrelated color.

In the present study, we aimed to better characterize the bi-directional influences between the VWM and oculomotor systems, here with a focus on automatic corrective saccades. Open questions still remain regarding this relationship. Whether task difficulty and an increased use of VWM differentially modulates oculomotor capture at the level of corrective saccades is unknown. The primary goal of our study was to investigate whether automatic corrective saccade biases depend on the strength of VWM representation. Using a modified version of Hollingworth & Luck’s (2009) paradigm, we quantified corrective saccade biases in response to different levels of task difficulty. Oculomotor capture as a function of varying task demands (in the form of different levels of memory load) has already been studied in a simple saccade task (Beck & Vickery, 2019), however here we manipulated task difficulty by varying the discriminability of the memorized target feature (color) from a foil in the memory test phase of the paradigm. Thus, lower discriminability should require a stronger, more precise representation of the memory color in VWM. This, we expect will be linked to a higher proportion of corrective saccade biases towards a memory color-matching distractor.

Our secondary question addressed the relationship between VWM and oculomotor control in the opposite direction. While (voluntary) saccade execution has been reported to influence VWM performance, do more automatic eye movements, such as corrective saccades and their biases, have a functional role by updating VWM content? Such an updating should lead to improved VWM performance after saccade corrections biases towards a distractor matching the memory color, and impair performance after correction biases towards a distractor of an unrelated color.

## 2. Methods

### 2.1. Participants

Twenty-seven participants were recruited among students of Phillips-Universität Marburg through the university’s online recruitment service, of which 20 participants’ datasets were kept for further analyses (see details on exclusions below). Participants were compensated with 8€/h or course credits upon completion of the experiment. All participants gave their informed consent in writing.

Inclusion criteria consisted of age between 18 and 40 years, being naïve as to the research question and hypotheses of the study, and normal or corrected-to-normal vision. Exclusion criteria consisted of color vision deficiency (verified prior to the experiment using the Ishihara test (Clark, 1924)), inability to execute stable eye fixations during the calibration of the eye tracker (1 excluded), inability to perform the two conditions of the VWM task within a given range of performance (40-80% accuracy in the ‘Difficult’ condition, 60-100% in the ‘Easy’ condition, verified offline separately for each participant prior to group analyses). If a participant’s average performance in the ‘Easy’ and ‘Difficult’ conditions did not differ by at least 10%, then we considered that our manipulation of the VWM load with the experimental conditions was unsuccessful for this participant and their data were excluded from further analyses (0 excluded for this reason). One participant was excluded because of technical reasons their eye tracking datafile was not saved after an experimental block. Eye movement data were inspected offline and trials categorized as “invalid”, following the criteria detailed in the next paragraph, were excluded. In case after data exclusion the number of remaining trials was below 50% of the total trial number, the participant’s data was excluded and replaced (5 excluded). We recruited until the usable sample reached 20 participants’ datasets.

Similarly to Hollingworth & Luck’s (2009) cleaning procedure, the majority of excluded trials were those where the gaze was not oriented towards the initial position of the saccade target, and where the eyes immediately fell on a colored disk instead of between two objects after array rotation were excluded (18.8% of total trials). Other, less frequent trial exclusions included trials where participants’ eye position did not fall within an invisible circle of a diameter of 1.3 dva around the center of the screen at the beginning of the fixation task (skipped trial) (3.2%), where eye position after the first (4.1%) or the second (corrective) saccade (2.8%) did not cross an invisible boundary of 75% of the distance between the center and the color disk array, where participants did not look at the target disk during the saccade task (1.7%), or where the disk array’s rotation occurred during a fixation (1.1%). 68.2% of the total trials were categorized as “valid” and used for further analyses.

The sample size was chosen based on the study of Hollingworth & Luck (2009), which our paradigm was based on, that included 16 participants per experiment. We reasoned that considering the slight modification of their paradigm, we increase the sample size to 20. This is well above sample sizes used in similar studies on oculomotor capture in relation to WM (ranging from 7 to 16 participants) (Soto et al., 2005, 2006; Olivers et al., 2006; Hollingworth & Luck, 2009; Hollingworth et al., 2013a; Ohl & Rolfs, 2018). Moreover, in our version of the task 100% of the trials included a rotation, as opposed to their 55%, meaning that our participants performed more trials in each of the three distractor color change conditions (192*1/3=64 trials per condition vs. their 264*0.55/3=48). This, coupled with the increased sample size, were implemented to assure higher statistical power.

All procedures were in accordance with the Declaration of Helsinki and were approved by the ethics committee of the Phillips-Universität Marburg, Department of Psychology (Proposal 2021-58k). The study was preregistered at https://doi.org/10.17605/OSF.IO/ZP9VX.

### 2.2. Equipment

The experiment was conducted using PsychoPy (v2022..2.5) (Peirce et al., 2019) and stimuli were presented on a VIEWPIXX monitor (VPixx Technologies Inc., Saint-Bruno, QC, Canada) with a size of 51.50 × 29 cm, a spatial resolution of 1,920 × 1,080 pixels and a refresh rate of 120 Hz. A viewing distance of 60 cm was maintained with the help of a chin and forehead rest.

Monocular eye movements were recorded from the eye with the better-quality signal during calibration, using an EyeLink 1000+ (SR Research Ltd., ON, Canada) at a sampling rate of 1,000 Hz.

### 2.3. Stimuli and procedure

Each participant ran the two conditions of the experiment (Easy and Difficult) in a semi-randomized order, so that the order of conditions was counterbalanced between participants. All stimuli were presented on a midlevel gray background (16.17 cd/m^2^). We used a modified version of the paradigm by Hollingworth & Luck (2009) consisting of a saccade task flanked by a visual working memory (VWM) task.

#### 2.3.1. VWM task

Each trial was initiated by the participant by simultaneously fixating a central fixation dot and pressing the space bar with their left hand. After the presentation of a white central fixation cross (height of 1 dva) for 350 ms, the memory stimulus, a square (side of 3.3 dva) whose color was selected semi-randomly from three categories (red, green, or blue), was presented for 300 ms. Within the selected color category, the particular value was selected from a set of 5 similar colors that were similarly spaced on the CIE Lab color coordinate system (see Supplementary Information for exact values and reasoning behind changing original color coordinates). With this spacing we sought to have sets of colors in which neighboring pairs have the same discriminability throughout the continuum of each color category. The memory stimulus was replaced by the fixation cross for 400 ms, then the Saccade task started (see next section) (Fig. 1).

**Figure 1.**
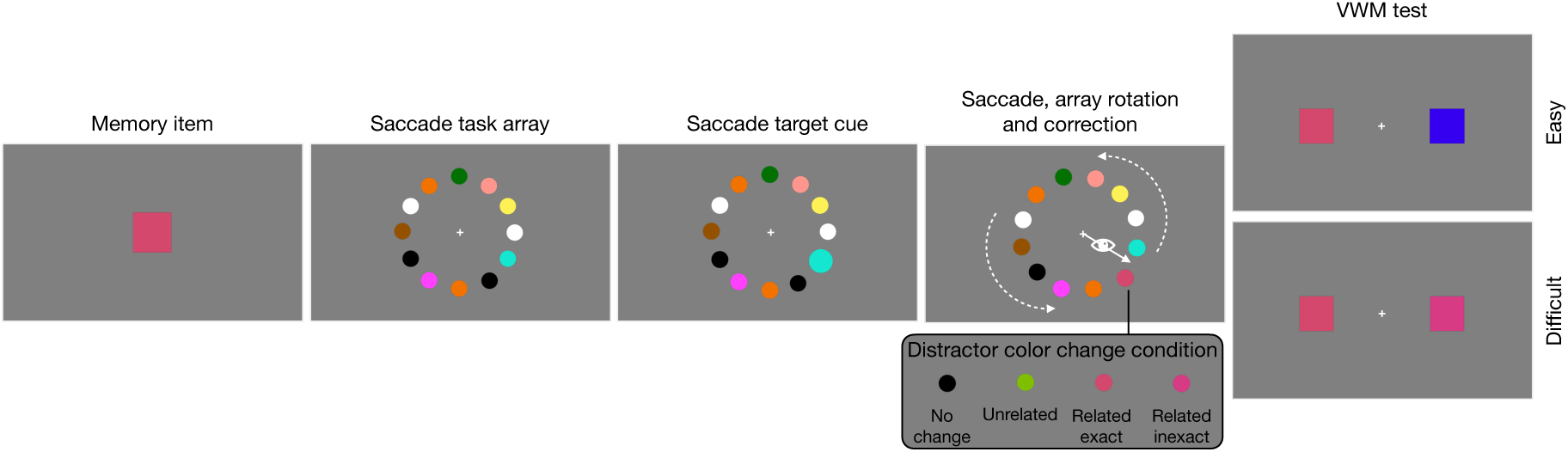
Illustration of the stimulus sequence in a trial. In this example, the black distractor changed color to the exact same color hue as the memorized one. The Related color change condition consisted of 50% Exact and 50% Inexact color changes. No, Unrelated, and Related change conditions were equiprobable and intermixed within each difficulty block. The white arrows indicating saccade direction and rotation direction are presented for illustration purposes and were not visible in the actual experiment.

The Saccade task was immediately followed by the test phase of the VWM task where two colored squares (side of 1.6 dva) were presented on the left and right side of the central fixation cross at a distance of 2.5 dva. Participants had to select the test stimulus that corresponded to the initial memory color via a left or right keypress. Task difficulty was manipulated in the two experimental conditions. In the Easy condition, the memory color had to be discriminated from a color belonging to a different color category (*e.g*., discriminating red and blue), thus other components of working memory could be recruited to perform the task (*e.g.*, semantic information of the category “red”). In the Difficult condition, the memory color had to be discriminated from a neighboring color from the same color category (*e.g.*, discriminating two perceptually similar red values). Thus, discrimination in this condition was likely exclusively based on VWM content (*i.e.*, no verbal color-categorization was possible). Incorrect answers to color discrimination received an auditory feedback in the form of a B flat tone played for 200 ms.

#### 2.3.2. Saccade task

The central fixation cross was surrounded by an array of 12 equidistant colored disks (diameter of 1.6 dva) located on an imaginary circle of a radius of 5.9 dva. The color of each disk in the array was selected semi-randomly from a set of 11 colors (excluding the color matching the current memory color category in a given trial) (see Supplementary Information for exact values). The same color could be present repeatedly in the array, however two disks of the same color were distanced by at least 2 other disks.

Participants were instructed to maintain central fixation. After a period of 1000 ms, the cue appeared, consisting of the rapid expansion for 50 ms then retraction of one of the circles. Participants then had to shift their gaze to the cued location.

Eye position was monitored and updated continuously. Saccade initiation was detected using the boundary technique with an invisible circle (diameter of 1 dva) whose position was updated at the moment the cue appeared. We concurrently checked that eye position was within an invisible circle of a diameter of 1.3 dva surrounding the central fixation cross. In order to make sure that the saccade to the target disk was always initiated from the center, if eye position was not within this imaginary boundary surrounding the center of the screen at the time of the cue, the trial was skipped, and a message reminded the participant to fixate the center, after which the program started the next trial. When eye position was detected outside the boundary, the position of the colored disk array was updated by an equiprobable clockwise or counterclockwise rotation of 15°. This manipulation allowed, at the end of the saccade, for the gaze to fall between two disks, the original saccade target and the distractor, inducing thus an artificial saccade error. After the saccade error was corrected towards one of the two disks, the array disappeared and was replaced by the central fixation cross for 400 ms before the test phase of the VWM task started. In case the gaze did not fall onto the target or distractor disks within 4 seconds a visual feedback was given in the form of a white square (side of 3.2 dva) appearing around the target disk for 300 ms. The program then proceeded to the memory test phase as usual.

Our experimental manipulation involved the correspondence between the content of VWM and the color of the distractor object. During the disk array’s rotation, while participants’ gaze was in-flight, the color of the distractor disk was manipulated in 3 equiprobable conditions that were intermixed within each difficulty block. In the ‘No change’ condition, the distractor’s color remained the same before and after the array rotation. In the ‘Unrelated change’ condition, the distractor’s color changed to a color from a different category than the one held in VWM. In the ‘Related change’ condition, the distractor’s color was changed to a value from the same color category as the memory color. Two equiprobable sub-conditions were used within the ‘Related change’ condition. In the ‘Exact change’ sub-condition the distractor color was the exact same value as the memory color from the VWM task. In the ‘Inexact change’ sub-condition the distractor color changed to a value neighboring the memory color.

In ‘Related’ and ‘Unrelated change’ trials the original disk array did not include the color from the same category (red, blue, or green) to which the distractor disk was changed.

### 2.4. Data analysis

Since two of our dependent variables (VWM performance and proportion of saccades towards target) were proportion data (i.e., percentages) bound between 0 and 1 and inappropriate for analyses of variance, we applied an arcsine square root transformation on these two measures before performing ANOVAs. This transformation deviated from our originally planned logit transformation described in our pre-registration since, while yielding similar transformed values, the arcsine square root transformation does not require making an arbitrary choice of a constant for correcting proportions of 0 and 1, which is necessary in the logit transformation. Note that in order to facilitate interpretation of the results, although analyses on proportion data were conducted on transformed data, we report back-transformed averages and confidence intervals, and the figures represent non-transformed data

### 2.5. Statistical analyses

All analyses were conducted using R (version 4.5.1, R Core Team, 2025) and the RStudio environment (version 2025.5.1.513, RStudio Team, 2016). As described in our pre-registration, we had a within-subject pseudo-randomized block design with 2 levels for the VWM load (*i.e.*, Easy *vs.* Difficult conditions) and 3 levels for the correspondence between VWM content and distractor color change (*i.e.*, No *vs*. Unrelated *vs.* Related change conditions). For sub-analyses in the Related condition only, the experimental design consisted of 2 levels for the VWM load and 2 levels for the distractor color change (i.e., Exact vs. Inexact). In the analyses of VWM performance an additional factor of direction of corrective saccade (i.e., towards target vs. distractor) was included. Each participant performed all conditions. Accordingly, we performed two-way repeated-measures ANOVAs and Bonferroni corrected pairwise t-tests to analyze our data (‘anova_test’ and ‘pairwise_t_test’ functions in the ‘rstatix’ package, Kassambara, 2023). Deviations from the sphericity assumption were corrected automatically via the Greenhouse-Geisser sphericity correction. Normal distribution of dependent variables was verified via the Shapiro-Wilk test for normality for each combination of factor levels. In case of non-normality, we performed non-parametric rank-based repeated-measures ANOVAs (‘rankFD’ package, Konietschke et al., 2022), followed by Tukey-adjusted pairwise comparisons using ‘mctp.rm’ function from the ‘nparcomp’ package (Konietschke et al., 2015), which applies ranked residuals to account for within-subject variability in multiple contrast testing. This function calculates an ANOVA-type statistic (ATS), which is a non-parametric equivalent of the F statistic. Alpha-error level was set to 0.05 throughout the analyses of the study. We report standard p-values and partial eta-squared values for effect size where applicable.

Additionally model fitting was used to verify (lack of) interaction effects. Bayesian mixed-effects logistic regression models were fitted to corrective saccade data using the ‘brms’ package (Bürkner, 2017) to predict the binary outcome of saccade correction of each trial (0=to distractor/incorrect, 1=to target/correct). Fitting was done using Bernoulli likelihood with logit link function, and weakly informative priors: Normal(0, 2.5) priors for regression coefficients, Student-t(3, 0, 2.5) prior for the intercept, and Student-t(3, 0, 2.5) prior for random-effect standard deviations. The performance of models with and without an interaction effect were evaluated using approximate leave-one-out cross-validation (LOO) and 10-fold cross-validation, using the *‘*loo’ package (Vehtari et al., 2017). Model comparison was done by computing the expected log predictive density (ELPD) and model weighting using the stacking method (Yao et al., 2018).

Binomial logistic mixed models were fitted to the performance data using the ‘lme4’ package (Bates et al., 2015) to predict participants’ binary responses on each trial (1=correct, 0=incorrect) with the three variables. Models containing different numbers of interaction effects were compared using Model Likelihood Ratio Tests in order to identify the most parsimonious model fitting the data. To assess potential multicollinearity among the predictors, adjusted Generalized Variance Inflation Factors (GVIF1/(2⋅df)) were calculated, since one of the categorical predictors had more than two levels. For task difficulty, GVIF1/(2⋅df)=1.00; for change condition, GVIF1/(2⋅df)=1.01; for saccade correction direction GVIF1/(2⋅df)=1.03. Given the low adjusted VIF values, multicollinearity among predictors was unlikely.

## 3. Results

### 3.1. Saccade correction bias

#### 3.1.1. Saccade correction accuracy

Saccade correction accuracy i.e., the proportion of corrective saccades directed to the target object after the artificial saccade error, was influenced by distractor change condition [ATS(1.96,101.78)=13.38, p<0.0001] as shown by a robust nonparametric repeated-measures ANOVA. Post hoc analyses showed that saccade correction accuracy, averaged over difficulty conditions, was lower in the Related (mean=0.85, 95% CI [0.78, 0.91]) compared to the Unrelated change condition (mean=0.95, 95% CI [0.93, 0.97], [p=1.88e-4]), which itself was lower than in the No change condition (mean=0.97, 95% CI [0.96, 0.99], [p=0.0098]), replicating the findings of Hollingworth & Luck (2009). Importantly, task difficulty also influenced saccade correction accuracy [ATS(1,101.78)=4.27, p=0.041], with lower average accuracy in the Difficult (mean=0.91, 95% CI [0.87, 0.94] than in the Easy condition (mean=0.96, 95% CI [0.94, 0.97]). No interaction between the two factors was found [ATS(1.96,101.78)=0.79, p=0.45] (Fig. 2A).

**Figure 2.**
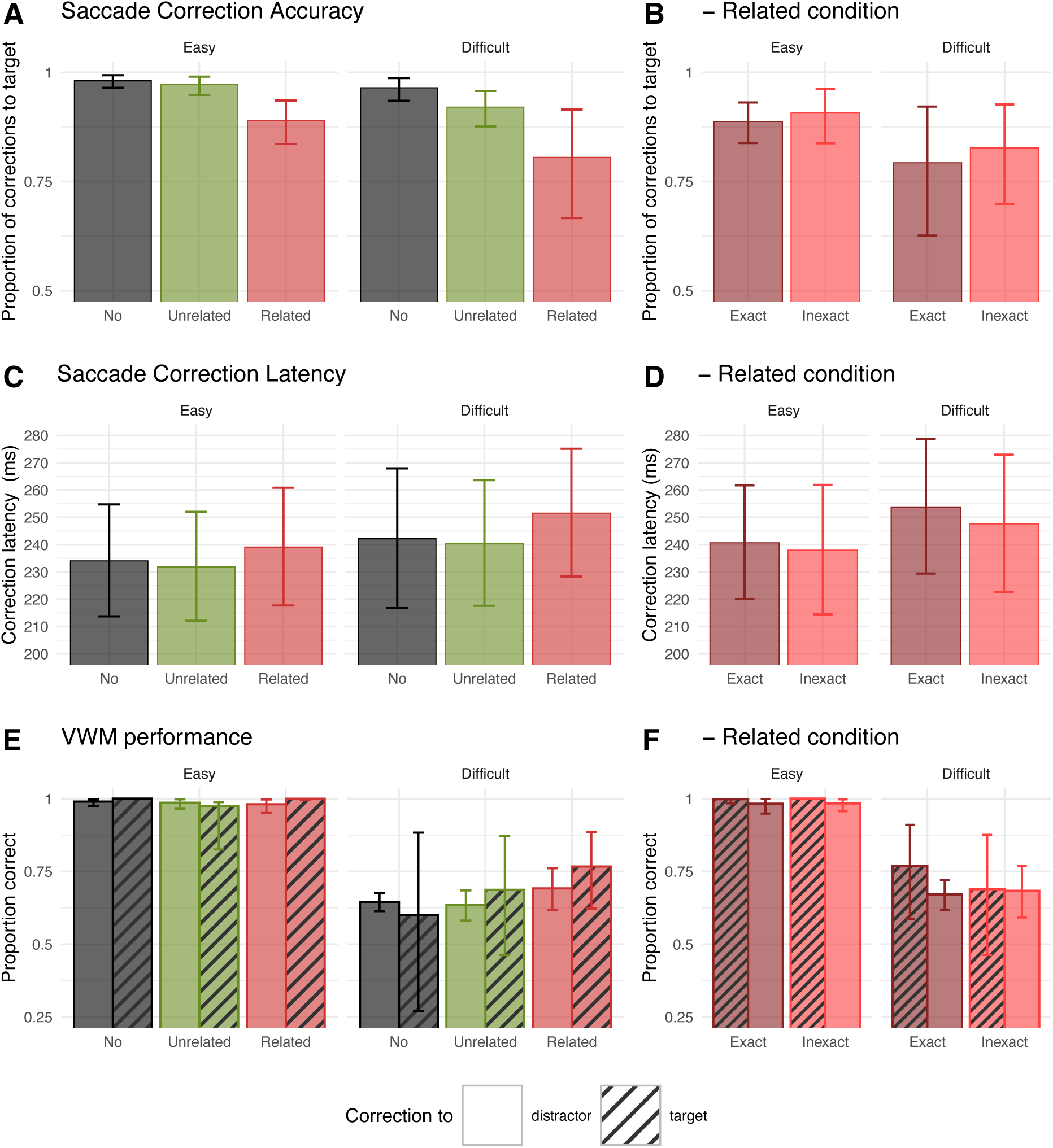
Main results. Average saccade correction accuracy, average latency of correct saccade corrections to the saccade target, and average performance in the VWM task by difficulty levels in the three distractor color change conditions (A, C, E) and in the two sub-conditions in the Related change condition only (B, D, F). In the graphs on VWM performance, empty bars correspond to trials where the saccade was corrected towards the saccade target and striped bars correspond to trials with corrections towards the distractor. Note that while statistical analyses on proportion data were conducted on arcsine square root transformed data, the figure represents back-transformed averages ±95% confidence intervals (error bars). See Supplementary Information for a version including individual data points.

As a means to verify an absence of interaction between our factors, we fitted two Bayesian logistic regression models predicting the binary outcome ‘saccade correction’ of each trial (0=to distractor/incorrect, 1=to target/correct): an additive model with task difficulty and change condition as predictors (correction ∼ difficulty + change condition + (1|subject)) and a model including their interaction (correction ∼ difficulty * change condition + (1|subject)). Subjects were included as random effect. Model comparison using leave-one-out cross-validation (LOO) indicated that the additive model provided a slightly better out-of-sample fit than the interaction model (difference in expected log pointwise predictive density, or elpd_diff = –1.9, SE = 0.9), a result replicated with 10-fold cross-validation (elpd_diff = –3.2, SE = 2.5). Stacking model weights further supported the additive model (weight_interaction_ ≈ 1.00 vs. weight_additive_<0.001), suggesting that including the interaction did not improve predictive performance. Inspection of the regression coefficients in the model with interaction confirmed this conclusion, as both interaction terms were centered near zero with narrow credible intervals: difficulty × condition1 (β = –0.03, 95%CI [–0.17, 0.11]) and difficulty × condition2 (β = 0.08, 95% CI [–0.10, 0.27]). Together, these results suggest that saccade correction accuracy is best explained by additive effects of task difficulty and change condition, without a meaningful interaction.

In the Related condition only, a repeated-measures ANOVA showed a tendential effect of task difficulty [F(1,19)=3.40, p=0.060, η_p_^2^=0.17] and a tendential effect of sub-condition [F(1,19)=4.34, p=0.051, η_p_^2^=0.19]. Although these effects did not reach statistical significance, numerically they matched the pattern of the previous analysis, with lower saccade correction accuracy in the Difficult (mean=0.81, 95% CI [0.71, 0.89]) than in the Easy condition (mean=0.90, 95% CI [0.86, 0.93]), and lower accuracy in the Exact condition, where the distractor matched exactly the color hue held in VWM (mean=0.84, 95% CI [0.77, 0.91]), than in the Inexact condition (mean=0.87, 95% CI [0.80, 0.93]). No interaction between the two factors was found [F=0.02, p=0.89, η_p_^2^=0.001] (Fig. 2B).

#### 3.1.2. Corrective saccade latency

Following Hollingworth & Luck’s (2009) study, we analyzed the latency of corrective saccades directed to the target object (excluding corrective saccades directed to the distractor). We excluded from this analysis trials where saccade latencies exceeded a cutoff value of ‘mean + 2 SD’ calculated for each difficulty level. The proportion of excluded trials was 4.1% in the Difficult and 3.5% in the Easy condition, similar to the proportion reported by Hollingworth & Luck (2009).

Corrective saccade latency was influenced by the distractor color change condition, as shown by a repeated-measures ANOVA [F(2,38)=3.86, p=0.030, η_p_^2^ =0.169]. Post hoc analyses reveled that the corrective saccade latency was higher in the Related condition (mean=246ms, 95% CI [230ms, 261ms]) than in the Unrelated condition (mean=236ms, 95% CI [222ms, 251ms], p=0.014). In the No change condition, the latency had an intermediate value (mean=238ms, 95% CI [223ms, 254ms]) and did not differ significantly from the other two conditions (No *vs.* Related p=0.24; No *vs*. Unrelated p=1.00). The effect of task difficulty was not significant [F(1,19)=2.02, p=0.171, η_p_^2^ =0.096], but average corrective saccade latency in the Difficult condition (mean=245ms, 95% CI [232ms, 258ms]) was numerically higher than in the Easy condition (mean=235ms, 95% CI [224ms, 246ms]) (Fig. 2C).

In the Related condition only, the effect of neither task difficulty, nor sub-condition, or the interaction between the two reached statistical significance, although numerically fitted the above-described pattern of results: latency was numerically higher in the Difficult (mean=251ms, 95% CI [234ms, 268ms]) than in the Easy condition (mean=240ms, 95% CI [224ms, 255ms]), and higher in the Exact color change (mean=247ms, 95% CI [232ms, 263ms]) than in the Inexact color change condition (mean=243ms, 95% CI [226ms, 260ms]) (Fig. 2D).

### 3.2. VWM performance as a function of saccade correction direction

Our secondary research question concerned the influence of the oculomotor system on VWM, specifically whether corrective saccade biases play a role in the updating of VWM content. This would be reflected by an interaction between saccadic correction direction and change condition.

A robust 3-way repeated-measures ANOVA confirmed that our experimental manipulation of task difficulty was successful, [ATS(1,82.63)=219.09, p<.0001], with higher average performance in the Easy (mean=0.99, 95% CI [0.98, 1.00]) compared to the Difficult condition (mean=0.67, 95% CI [0.61, 0.73]). Saccade correction direction also influenced VWM performance [ATS(1,82.63)=11.61, p=.0010], with higher average performance in trials with corrections to the distractor (mean=0.89, 95% CI [0.83, 0.94]) than to the target (mean=0.87, 95% CI [0.83, 0.90]). Nonetheless, no other effects were found: the direction of saccade correction did not interact with task difficulty or change condition (Fig. 2E).

The difference in performance between the two difficulty blocks was considerably larger than that of the change conditions. In order to verify that this strong effect of difficulty did not overpower more subtle interaction effects within difficulty blocks, we chose to conduct exploratory sub-analyses. In the Easy condition, a robust 2-way repeated-measures ANOVA revealed a main effect of saccade correction direction [ATS(1,69)=19.27, p<.0001] with higher average performance following saccade correction to the distractor (mean=1.00, 95% CI [0.98, 1.00]) than following saccade corrections to the target (mean=0.99, 95% CI [0.97, 0.99]). The effect of condition or the interaction between the two factors were not statistically significant. In the Difficult condition, according to a similar robust 2-way repeated-measures ANOVA, neither the effect of saccade correction direction or that of change condition, nor the interaction between the two factors reached statistical significance. The result of these sub-analyses suggest that the main effect of correction direction in the 3-way ANOVA was most likely driven by the Easy condition, as the effect was absent in the Difficult condition.

As an additional verification of the absence of interaction between change condition and saccade correction direction, we fitted a logistic mixed model to the data in order to relate the binary outcome ‘response’ of each trial (0=incorrect, 1=correct) to the three factors as predictors (response ∼ difficulty + change condition + correction direction + (1|subject)). Subjects were included as random effect. Next, we compared this model to one containing a double interaction between factors (response ∼ difficulty * change condition * correction direction + (1|subject)) and one containing a single interaction between the critical factors in question (response ∼ difficulty + change condition * correction direction + (1|subject)). Model comparisons showed that the fit of the model was not significantly improved by the addition of a double interaction (AIC_no-interaction_=4087.3, AIC_double-interaction_=4090.0, χ^2^=11.38, p=0.12) or a single interaction (AIC_single-interaction_=4091.3, χ^2^=0.03, p=0.98), meaning that the more parsimonious (*i.e.*, simpler) model fits our data better.

In the Related condition only, a separate robust 2-way repeated-measures ANOVA revealed a main effect of task difficulty [ATS(1,61.88)=148.87, p<.0001] with, again, higher average performance in the Easy (mean=0.99, 95% CI [0.99, 1.00]) compared to the Difficulty condition (mean=0.70, 95% CI [0.63, 0.76]), as well as a main effect of saccade correction direction [ATS(1,61.88)=6.66, p=.012], with higher performance following saccade correction to the distractor (mean=0.93, 95% CI [0.87, 0.97]) than following saccades to the target (mean=0.87, 95% CI [0.83, 0.91]). Again, no interaction between sub-condition and correction direction, or any other statistically significant effect was found (Fig. 2F).

## 4. Discussion

The present study examined the bi-directional link between VWM and corrective saccades through two main experimental question.

### 4.1. The influence of VWM content on corrective saccade biases

Our first question concerned whether oculomotor capture is dependent on the strength of VWM representation. This was operationalized by manipulating the difficulty of our VWM task in separate blocks. Our rationale was that increasing task difficulty, by lowering the discriminability of the target color from the foil in the test phase, engages the visual component of WM to a higher extent compared to a condition where target and foil colors are easily discriminable and non-visual strategies, *e.g.*, memorizing verbal cues, can be used.

First, we verified that our experimental manipulation of task difficulty was successful, as shown by clear differences is VWM performance between the Difficult and Easy blocks throughout the analyses of performance. All participants’ performances fell within the predefined, preregistered performance ranges, showing that our manipulation of color hue discriminability in the two conditions was robust. Our recreation of a more difficult and an easier version of Hollingworth & Luck’s experiment was successful, as average performance levels were respectively lower (0.67) and higher (0.99) than that in the original study (0.718).

Second, the proportion and latencies of saccade corrections towards the saccade target varied depending on the color the distractor changed to. When the distractor changed to a color that matched the memorized one (Related condition), saccades were more biased towards the distractor (and away from the original saccade target) than in a condition where the distractor changed to a color that was unrelated to the one held in memory. The effect was also apparent in saccade latencies, with slower gaze corrections to the saccade target when it was neighboring a distractor that matched the memory color. Note that the difference between Unrelated and No conditions in VWM performance points to a general attentional effect related to the color change itself. The critical effect related to our question is the difference between the Unrelated and Related conditions. Our results replicate the findings of (Hollingworth & Luck, 2009) in an easier and a more difficult version of their task, corroborating that the content of VWM can influence oculomotor control.

Thirdly, and most importantly for our research question, the proportion of saccade correction bias was higher in the Difficult compared to the Easy condition, but difficulty did not interact with distractor color change conditions. This lack of interaction, confirmed by both frequentist and Bayesian analyses, suggests that difficulty has a general effect on corrective saccade biases. Increased difficulty seems to amplify corrective saccade biases overall, but not modulating the previously described effect of distractor color change. This finding can be interpreted in light of previously proposed effects of increased distractor interference in high working memory load conditions (Lavie et al., 2004; Lavie & De Fockert, 2005; Konstantinou et al., 2014). According to the proposal, loading working memory in a secondary task uses cognitive control resources that are shared with selective attention, leading to stronger interference from task-irrelevant distractors. In our experiment, the Difficult condition requires the exact representation of the target color hue in VWM for its discrimination from another similar hue, and the use of other strategies in discriminating the two hues, e.g. verbally, is unlikely. This substantial increase in visual WM load can compete with cognitive control resources in our secondary saccade task and facilitate distractor interference, as reflected by higher oculomotor capture at the saccade correction level (*i.e.*, corrective saccade bias).

Our findings on the effect of task difficulty on saccade correction biases are in contrast with findings on attentional capture by Olivers et al. (2006). In their study distractors matching or sharing features with an object held in WM interfered with a visual search performance (as reflected by slower response times) only when the memory content was difficult to verbalize and when participants were encouraged to use “a more visual kind of memory” rather than memorizing verbal cues which lead to no interference with search performance. It appears that, as opposed to attentional capture measured via search times in Olivers et al.’s study, oculomotor capture at the level of automatic corrective saccades permeates difficulty levels in our task and is present independently of the type of representation activated in WM: both when the exact representation of the target color hue has to be maintained in VWM (our Difficult condition) and when other components of WM (*e.g.*, verbal) can be used (our Easy condition). This suggests that the memory target color hue is represented at a lower level than that playing a role in the attentional capture effects described above, remaining active independently of task difficulty and biasing automatic saccadic eye movements. Our results thus point to a dissociation between attentional capture effects and oculomotor capture, at least at the automatic level of corrective saccades.

It is noteworthy that Olivers et al. also measured oculomotor capture effects in their Exeriment 7 (in parallel with their attentional capture effects), however they observed it in a single difficult condition where participants were instructed to use visual WM and they did not manipulate task difficulty.

### 4.2. The influence of corrective saccade biases on VWM performance

Our secondary research question addressed whether saccade biases during automatic saccade corrections have a functional role in updating VWM content. Making an additional eye movement to a feature that is critical to the VWM task at hand could potentially refresh VWM content and improve performance. Conversely, making an eye movement to a feature that does not match the memory target, refreshing to the new feature would impair subsequent performance. Such an influence of eye movements on VWM would thus close the circle of bi-directional influences between VWM and the oculomotor system. However, our results revealed no strong evidence of such an influence of corrective saccades on VWM performance. As shown by the frequentist analyses (main and sub-analyses) and verified with the model comparison method, there was no interaction between corrective saccade direction and distractor change condition that would support an updating of VWM content by saccade biases.

A main effect of corrective saccade direction was, however, significant in the main analysis of VWM performance. Sub-analyses showed that this was driven by the Easy condition with higher VWM performance following corrections to distractors than saccade targets, although the difference was small and performance in general in this condition was close to perfect (on average 97% correct responses). It is important to note that the Easy condition contained intrinsically fewer trials with saccade corrections to the distractor than the Difficult condition, and the imbalanced numbers between ‘saccade target’ and ‘distractor’ trials calls for caution when interpreting this effect. The small number of ‘distractor’ trials means that a ceiling effect can easily be reached and distort an actual effect of these trials. In the Difficult condition, where more corrections to the distractor occur and the ratio of ‘saccade target’ to ‘distractor’ trials is more balanced, the effect of corrective saccade direction on VWM performance was absent. In all, we found no evidence of corrective saccade biases having a role in the updating of VWM content.

### 4.3. Attentional selection may be necessary for the saccade system to influence VWM

What conditions are required for the oculomotor system to have an influence (in a beneficial or detrimental way) on VWM? Several studies addressed this relationship. Tas et al. (2016) found that selecting and making a saccade to a secondary object, that is unrelated to the one held in VWM, interfered with VWM performance, while covertly attending to it (*i.e.*, shifting attention to the object without making a saccade) does not interfere with VWM. Their interpretation was that executing a saccade leads to the automatic encoding of the saccade target in memory and its representation will compete for VWM resources, whereas covert attention leads to no such encoding of the target. Hamblin-Frohman & Becker (2019) challenged this interpretation by proposing that covert attention may require only spatial attention which would not interfere with VWM. In contrast, overt attentional orienting requires object features to be processed to a higher level, and feature based attentional selection would share resources with VWM (if not equate it). They indeed showed that attending to an object feature, with or without saccading to it, was enough to interfere with VWM performance. This suggests that, in fact, feature based attentional selection, but not spatial attention, shares resources with VWM (Hamblin-Frohman & Becker, 2023).

Importantly, our results (specifically those of the Difficult condition with higher statistical power) indicate no interference with VWM performance when VWM related saccade biases are executed. A possible reason for this may be that feature based attentional selection may be limited to voluntary saccades and is not involved in the execution of corrective saccades which are a more automatic form of eye movements. This is consistent with the findings of Schut et al. (2017) that, while performing an additional saccade during a VWM task does impair VWM performance compared to a “No saccade” condition, executing a corrective saccade after the voluntary saccade yields no additional interference with performance. While this corroborates our results, a critical difference between Schut et al.’s paradigm and ours is that, in their study the corrective saccade is made to an item completely unrelated to the memory item. In our experiment, however, the saccade target sometimes matches the critical feature (*i.e.*, color) of the memory item. It is remarkable that looking at the feature that is critical to the VWM task at hand a second time does not yield a clear benefit to performance.

It is all the more surprising that it has been shown that merely making an eye movement to a cued location shortly *after* the disappearance of an array of items increases the strength of the VWM representation of the item that was previously at that location (Ohl & Rolfs, 2017, 2018; Hanning & Deubel, 2018). This effect is most likely driven by oculomotor and pre-saccadic attentional selection, as the WM performance benefit is present when the saccade is executed to the item, but also when the saccade to the selected item is only prepared, but is never executed (Hanning et al., 2016).

In light of these studies, attentional selection related to saccade execution may be a necessary condition for the saccade system to influence VWM content. Attentional selection, however, may be limited to voluntary saccades, meaning that more automatic eye movements, like corrective saccades (and their biases towards distractors) may not have bi-directional links with the VWM system. Instead, they would bypass the connection between oculomotor control and VWM systems that is supported by (feature based) attentional selection, and take a more automatic, likely non-attentional pathway which may be uni-directional (WM ➞ eye movements). This pathway may support the primary function of corrective saccades, *i.e.*, stabilizing an erroneous saccade and bringing the gaze onto the item stored in VWM. It would also allow VWM to bias involuntary, automatic eye movements in a uni-directional manner toward items that match VWM content. Such a lower-level pathway, that bypasses feature based attentional selection mechanisms, would complement a higher-level network between VWM and the (voluntary) saccade system, in which bi-directional links have been described (van Ede, 2020), and underlie automatic eye movements in the complex neuronal system of action and perception.

## Acknowledgments

We thank Marten Stegers for his help with data collection. The study was supported by the Deutsche Forschungsgemeinschaft (DFG, German Research Foundation) – project number 222641018 – SFB/TRR 135 TP B2.

