## Supplementary Information for "Visual working memory-related saccade biases are amplified by task demands, without updating working memory content"

Patrik Polgári & Alexander C. Schütz
Sensorimotor Learning Unit, Department of Psychology, University of Marburg, Germany

**Redefinition of color coordinates:**

Hollingworth & Luck (2009)'s original paradigm used similar memory and foil color hues in the test phase of the VWM task. When trying to recreate the paradigm as our "Difficult condition" with the colors used in the original study (defined in the 1931 CIE color coordinate system), some color pairs were easier to distinguish than others, making the task's difficulty uneven throughout trials. Thus, we chose to use new sets of color coordinates such that their coordinates in the CIE lab coordinate system are equidistant within a color set (*e.g.*, 5 blue values), making them perceptually equidistant. To achieve an even difficulty between trials, in the memory test phase of the Difficult condition we always used pairs of colors that were neighboring each other within a color set (*e.g.*, "red2" and "red3"). In the Easy condition, a strict control of the choice of exact color values was not necessary, since colors from different categories (*e.g.*, a blue and a red) were presented in the test phase.

**List of CIELab coordinates of the colors used as memory targets/foils in the VWM task:**

'red1': [50.0, 72.0, 20.0],
'red2': [50.0, 70.0, 22.0],
'red3': [50.0, 67.0, 33.0],
'red4': [50.0, 64.0, 40.0],

'red5': [50.0, 57.0, 46.0],

'blue1': [37.7, 14.2, -74.7],

'blue2': [37.7, 19.2, -72.7],

'blue3': [37.7, 24.2, -70.7],

'blue4': [37.7, 32.2, -68.7],

'blue5': [37.7, 40.2, -64.7],

'green1': [64.3, -72.0, 26.5],

'green2': [64.3, -68.0, 31.5],

'green3': [64.3, -64.0, 39.5],

'green4': [64.3, -58.1, 44.5],

'green5': [64.3, -52.1, 52.5]

**List of CIELab coordinates of the colors used in the stimulus array in the Fixation task:**

'red': [48.134, 76.002, 68.356],

'blue': [38.551, 59.762, -100.418],

'green': [38.725, -43.373, 45.623],

'yellow': [91.819, -18.197, 97.355],

'magenta': [60.782, 93.143, -66.044],

'black': [0.005, -0.019, 0.008],

'white': [100.000, 0.000, -0.000],

'brown': [38.021, 11.268, 33.511],

'pink': [66.749, 41.524, 8.617],

'orange': [59.798, 45.019, 71.724],

'aqua': [87.969, -41.583, -20.914]

**Figure: Main results with individual datapoints.**

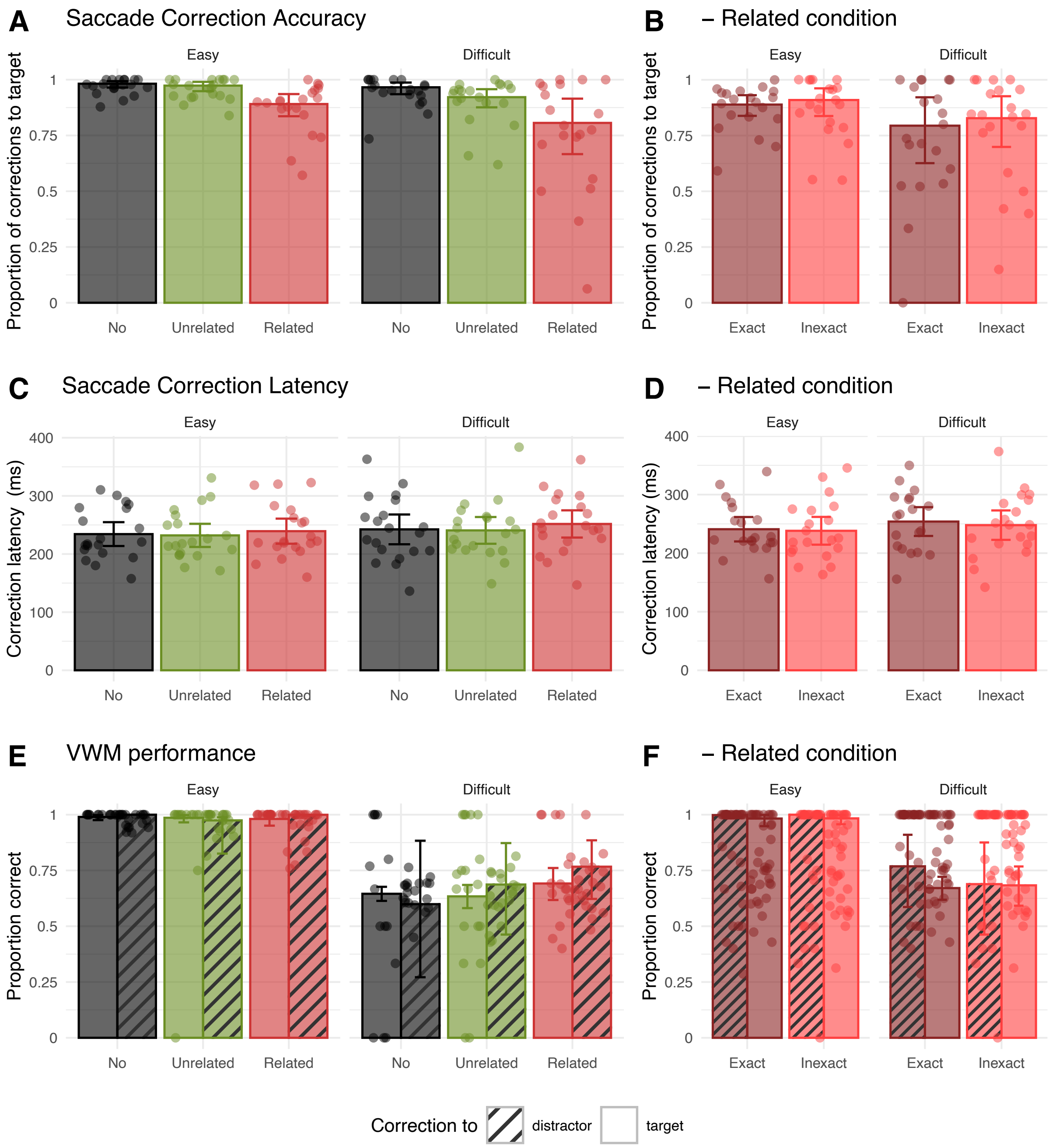


Average saccade correction accuracy, average latency of correct saccade corrections to the saccade target, and average performance in the VWM task by difficulty levels in the three distractor color change conditions (A, C, E) and in the two sub-conditions in the Related change condition only (B, D, F). In the graphs on VWM performance, empty bars correspond to trials where the saccade was corrected towards the saccade target and striped bars correspond to trials with corrections towards the distractor. Note that while statistical analyses on proportion data were conducted on arcsine square root transformed data, the figure represents back-transformed averages ±95%

confidence intervals (error bars), and non-transformed individual data points (circles). Note that the y-axes have a different scales here than in the main manuscript.
